# How NINJ1 mediates plasma membrane rupture and why NINJ2 cannot

**DOI:** 10.1101/2023.05.31.543175

**Authors:** Bibekananda Sahoo, Zongjun Mou, Wei Liu, George Dubyak, Xinghong Dai

## Abstract

NINJ1 is a recently identified active executioner of plasma membrane rupture (PMR), a process previously thought to be a passive osmotic lysis event in lytic cell death. NINJ2 is a close paralog of NINJ1 but it failed to mediate PMR. By cryoEM, we found that both NINJ1 and NINJ2 were able to assemble into linear filament that binds strongly to lipids on one side but is water-soluble on the other side. The more-or-less straight NINJ1 filament was able to wrap around a membrane bleb and solubilize it from the plasma membrane to induce PMR; however, the intrinsically curved NINJ2 filament failed to do so, explaining its incapability of mediating PMR. We further demonstrated that binding to cholesterol at the inner leaflet of the lipid bilayer was responsible for the curving of the NINJ2 filament, while strong lipid binding at the outer leaflet was contributing to NINJ1’s capability of mediating PMR.

## Introduction

Lytic cell death is an inflammatory cell death caused primarily by pyroptosis and necroptosis^1,2^. Plasma membrane rupture (PMR) is a cataclysmic event at the end of lytic cell death that releases large-size intracellular molecules known as damage-associated molecular patterns (DAMPs), which in turn amplify the inflammatory response^3,4^. In contrast, non-lytic cell death, mainly referring to apoptosis, does not leak intracellular components and trigger inflammatory responses. Lytic cell death plays an important role in host defense against pathogen infections, but its dysregulation is also implicated in many inflammatory diseases and pathological conditions^5^. PMR was long thought to be a passive event caused by osmotic swelling of the cell after plasma membrane leakage induced by gasdermin-D (GSDMD) pores in pyroptosis^6–11^ or the mixed-lineage kinase domain-like protein (MLKL) oligomers in necroptosis^12–17^. Until recently, a study identified ninjurin-1 (NINJ1) to be an active executioner of PMR^3^. NINJ1 is a 16-kDa plasma membrane protein previously identified as a nerve injury-induced protein (hence the name ninjurin) whose upregulation in injured neurons and in surrounding Schwann cells was thought to promote axonal regeneration through homotypic adhesion^18–20^. NINJ1 was predicted to have two transmembrane helices and one extracellular amphipathic helix. It undergoes oligomerization to induce PMR, and the amphipathic helix was shown to play a key role in this process^3^. However, a highly homologous protein NINJ2 in the plasma membrane bearing a similar amphipathic helix could not induce PMR^3,21^, further obscuring a mechanistic understanding of NINJ1-mediated PMR. To understand how NINJ1 mediates PMR and why NINJ2 cannot, we use cryogenic electron microscopy (cryoEM) to determine and compare the structures of NINJ1 and NINJ2.

## Results

### NINJ2 and NINJ1 both form linear oligomers

Human NINJ1 and NINJ2 share 52% sequence identity, yet NINJ2 is not capable of mediating PMR, nor showing cytotoxicity when overexpressed in HEK293T cells, as NINJ1 is^3^. Since it has been demonstrated that the induction of PMR is correlated with NINJ1 oligomerization^3^, we wanted to test if NINJ2 is capable of oligomerization or not. We transiently expressed N-terminally 3xFLAG-tagged human NINJ2 in suspension Expi293F cells, and purified it from the membrane fraction with detergent solubilization followed by peptidisc reconstitution^22^. A very broad peak was observed in size-exclusion chromatography (SEC) (FigureS1A), indicating largely varying sizes of the molecules. Indeed, cryoEM of three representative fractions showed a spectrum of oval- to belt-shaped particles. The 2D class averages indicated that they were linear assemblies of a helix-rich subunit at various lengths (FigureS1B). A preliminary trial of human NINJ1 overexpression and purification following the same strategy generated a similar SEC profile, albeit at much lower yield due to severe cytotoxicity^3^, and the particles looked similar to those of NINJ2 under negative-staining EM (data not shown). In summary, in vitro purified NINJ1 and NINJ2 form linear oligomers in a similar fashion.

### CryoEM structure of the NINJ2 filament

To understand the molecular interactions responsible for NINJ1 or NINJ2 oligomerization, we sought to determine atomic structures of those oligomers using cryoEM. First, we tried different strategies of membrane protein purification for NINJ2 and found that exchange the detergent with amphipol using the stepwise dilution method (see detail in Materials and Methods) helped to generate relatively long and uniform NINJ2 oligomers, as indicated by a sharp peak close to the void volume in the SEC profile (FigureS2A). Negative-staining EM and cryoEM showed both straight and slightly curved thin filaments in the length of 50-70 nm (FigureS2B and S2C). CryoEM 2D class averages suggested that the straight ones were the top or bottom views of the filament if we define the curved ones as the side views (Figure1A). Initial cryoEM data processing indicated that the particles had preferred orientations. By collecting more data with the sample tilted to 30°, and carefully balancing the number of particles from different views to be included for 3D reconstruction, we managed to obtain a structure of the NINJ2 filament at 3.07 Å resolution (Figure1B-E, FigureS3A-D), which enabled us to build a reliable atomic model (Figure2B, TableS1).

**Figure 1.**
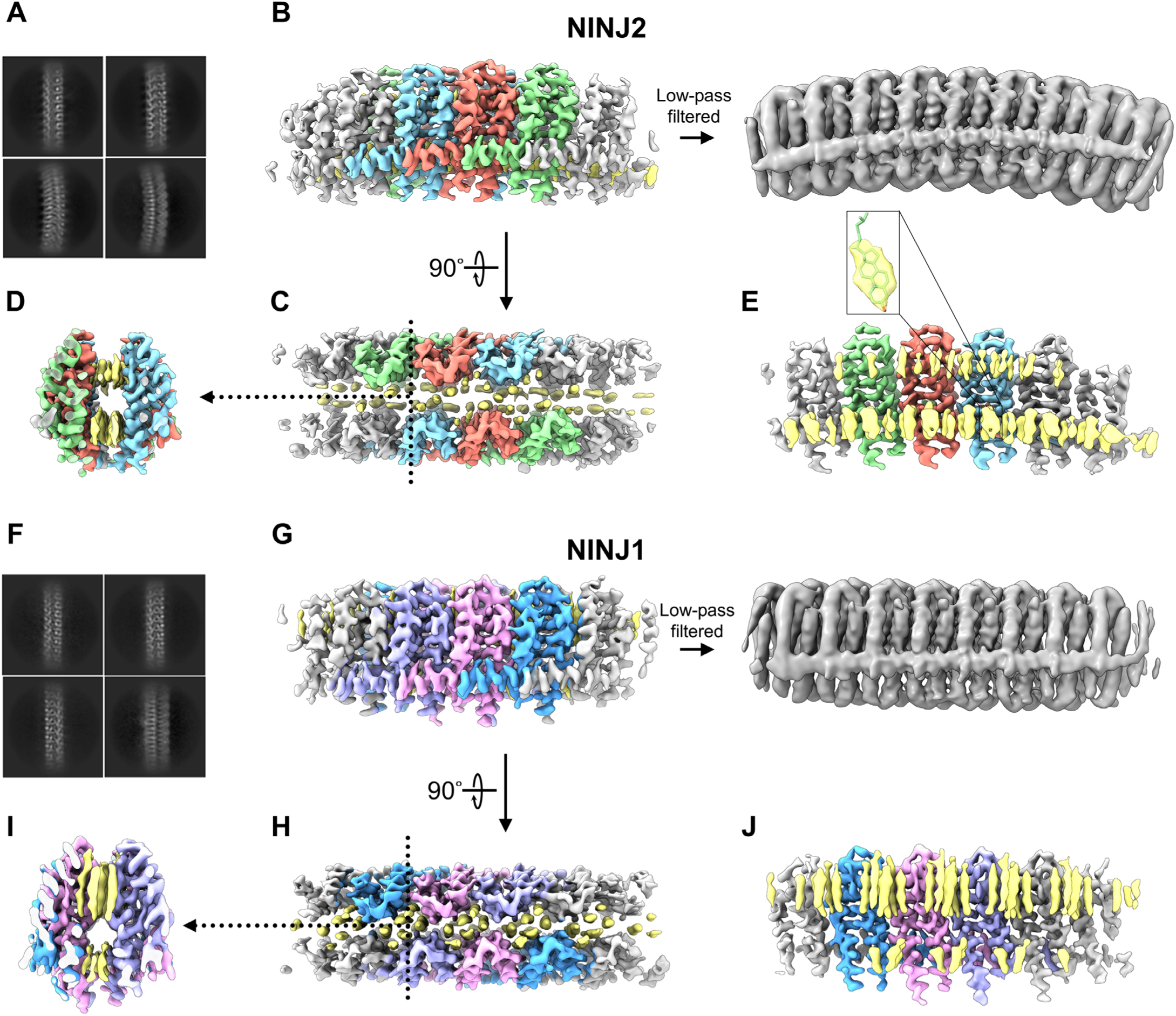
CryoEM Structures of the NINJ2 and NINJ1 Filaments. (A) Representative 2D class averages of cryoEM images of the NINJ2 filament. (B-E) Side view (B), top view (C) and clipped views (D, E) of the cryoEM density map of the NINJ2 filament. Three subunits in each protofilament are colored, rest as grey. Lipid densities are colored as yellow. Right panel in (B) shows a low-pass filtered version of the map to better illustrate the curved shape of the NINJ2 filament. Dotted line in (C) marks the clipping plane for the view shown in (D). One of the two protofilaments was removed in (E) to expose the lipid densities. The magnified inset in (E) highlights one of the cholesterol-like densities. (F) Representative 2D class averages of cryoEM images of the NINJ1 filament. (G-J) Side view (G), top view (H) and clipped views (I, J) of the cryoEM density map of the NINJ1 filament, following the same displaying scheme as explained for NINJ2.

The NINJ2 filament looked like a slightly arched bridge, with two sidewalls built up by interlinked NINJ2 proteins running in anti-parallel directions, sandwiching two layers of lipid like the deck of the bridge (Figure1B and 1C). Geometry of the two lipid layers obviously resembled the bilayer of a biological membrane (Figure1D and 1E), suggesting that the filamentous structure was likely formed in the cell membrane—and thus was biologically relevant—instead of being triggered by detergent solubilization. Of particular interest, a regular pattern of cholesterol-like densities with the very characteristic planar steroid ring was identified in the inner leaflet of the bilayer (Figure1E). The other lipid densities were not so well resolved (Figure1E), suggesting that they were less specific as compared to the cholesterol in associating with the NINJ2 protein.

### CryoEM structure of the NINJ1 filament

We applied the same strategy of detergent solubilization followed by amphipol exchange to purify NINJ1 oligomers for cryoEM structure determination. Our initial trials always ended up with very low yield when we used the cell pellet for purification as we did for NINJ2. However, when we turned to the supernatant after cell pelleting, and solubilized the small debris collected from there by ultracentrifugation, the yield of NINJ1 boosted to a similar level as that of NINJ2 purified from the cell bodies (FigureS2D). This observation suggested that most of the NINJ1 proteins were released to the extracellular space, likely in association with small membrane blebs, as a consequence of NINJ1-mediated PMR.

The purified NINJ1 oligomers showed a similar SEC profile to that of NINJ2 (FigureS2D). The particles also looked similar to those of NINJ2 under negative-staining EM and cryoEM with filamentous shape, but they appeared to be slightly skinnier and less curved compared to the NINJ2 filaments (FigureS2E and S2F). The differences became more prominent when cryoEM 2D class averages were compared (Figure1F and 1A). Particularly, the side view of NINJ1 filament was straight (Figure1F), in stark contrast to the curved side view of NINJ2 (Figure1A). Structure determination of the NINJ1 filament was done in a similar way as NINJ2, with tilted images combined with non-tilted ones to cope with the preferred-orientation problem. Resolution of the final density map was 2.75 Å (FigureS3E-H).

As expected from observations in 2D, the NINJ1 filament was nearly straight (Figure1G), in contrast to the curved NINJ2 filament (Figure1B). Yet it had the same architecture as NINJ2, with two anti-parallel proteinaceous sidewalls sandwiching a thin lipid bilayer (Figure1H and 1I). In addition to the different curvatures, another notable difference between the two filaments was in their associated lipids, particularly those at the outer leaflet of the bilayer (Figure1J and 1E). Compared to the relatively weak but continuous densities in NINJ2 (Figure1E and Figure4G), the outer leaflet densities in the NINJ1 filament were strikingly stronger and in a punctate distribution, with six fatty acid chains clustered in each punctum (Figure1J and Figure4A). Each of these fatty acid chain densities in NINJ1 could find a match in NINJ2 at the corresponding position relative to the protein sidewalls. It could be regarded as if one central layer of lipids present in the NINJ2 filament were missing in the NINJ1 filament (Figure4A and 4G). This explained the above observation in 2D that the NINJ1 filament appeared to be slightly skinnier than the NINJ2 filament. The inner leaflet lipid densities in NINJ1 also appeared to be discrete as compared to those in NINJ2 (Figure1J and 1E, Figure4B and 4H). These differences in lipid binding between NINJ1 and NINJ2 turned out to have important implications in their different capabilities of mediating PMR as discussed later.

### Atomic interactions for the assembly of the NINJ1 and NINJ2 filaments

Despite the notable difference in the global filament structure as described above, the individual subunits of NINJ1 and NINJ2 were highly similar (Figure2A and 2B, FigureS7A), and their atomic interactions leading to the assembly of the filaments were highly conserved too. Human NINJ2 has 142 amino acids (a.a.), while NINJ1 has 152 a.a., with a major insertion near the N-terminus (FigureS4). However, for the regions that were well resolved in our structures, from Asn39 to Phe139 for NINJ1 and Asn25 to Phe125 for NINJ2 (Figure2A and 2B), every residue could be matched from one to the other (FigureS4) and each pair were located at roughly the same position, even for those in the loop regions (FigureS7A). The root mean square deviation (RMSD) for the 101 pairs of C-alpha atoms was measured to be 0.792 Å. For clarity, in the following descriptions, NINJ1 sequence would be used, with the corresponding NINJ2 sequence given in parenthesis. Both NINJ1 and NINJ2 were predicted to have a flexible, extracellular N-terminal coil, an amphipathic helix (AH), followed immediately by two transmembrane helices (named TM1 and TM2 here), and an extracellular C-terminus. The Alphafold2 predicted structures^23^ also bear three helices. However, in both NINJ1 and NINJ2 filaments, the subunit was a four-helix structure; the predicted single amphipathic helix had divided into two short ones, named AH1 and AH2 here, with a nearly orthogonal kink at residue Leu52 (Leu38) (Figure2A and 2B). AH2 was bundled with the transmembrane helices, with its hydrophobic sidechains of Leu52 (Leu38), Ile54 (Val40), Leu57 (Phe43), Ala61 (Ala47), Leu64 (Leu50), Val67 (Val53) and Val68 (Leu54) inserted into a highly hydrophobic groove formed by TM1 of the same protomer and TM2 of a neighboring protomer (Figure2C and 2G). In addition to those hydrophobic interactions, Asn60 (Asn46) on AH2 seemed to be important as it formed a network of hydrogen bonds, likely mediated by a H_2_O molecule, with a highly conserved triad of polar residues, Ser87 (Ser73), Gln91 (Gln77) on TM1 and Asn133 (Asn119) on TM2, all from the same protomer as Asn60 (Asn46) (Figure2D and 2H).

**Figure 2.**
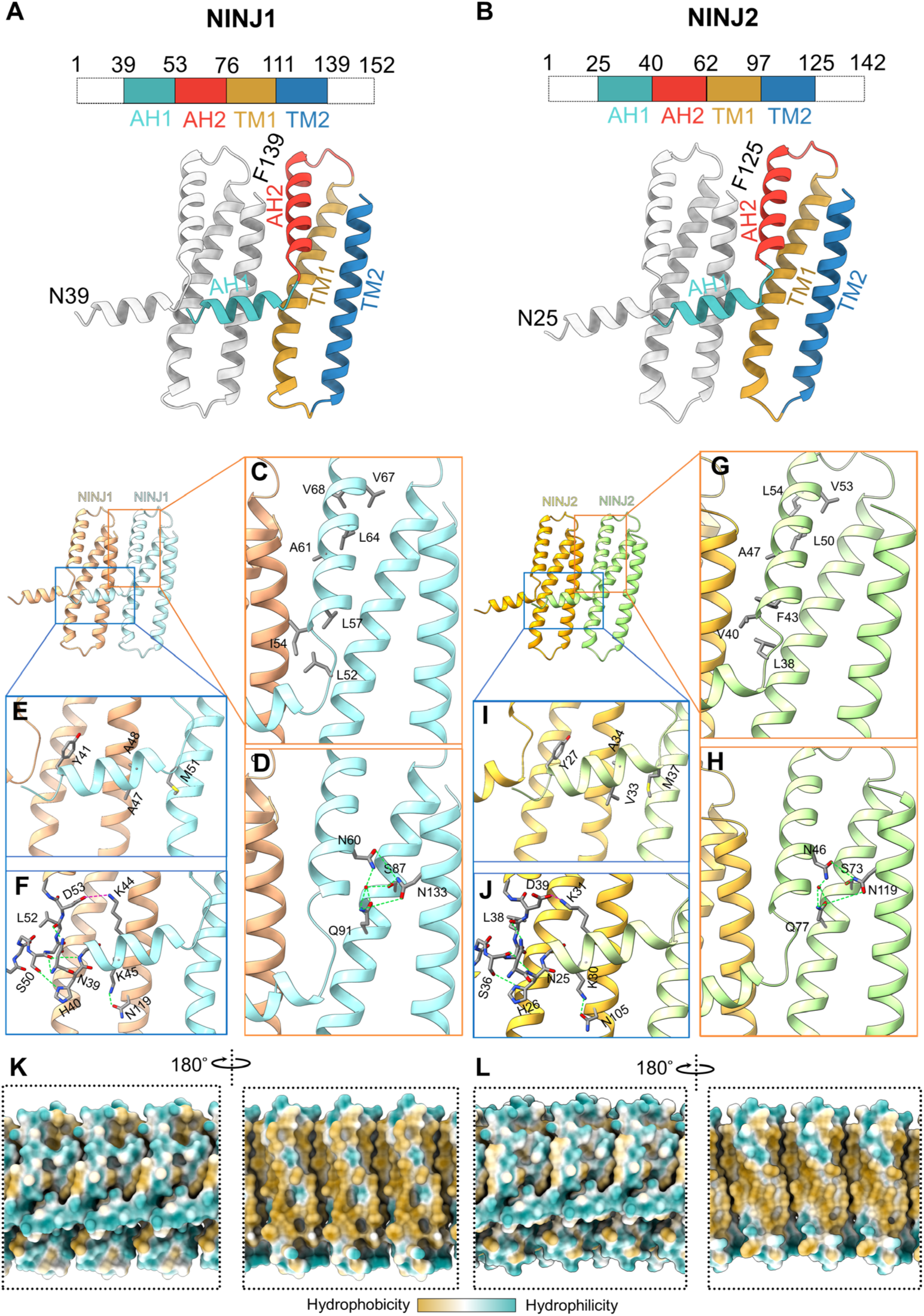
Atomic interactions for the assembly of the NINJ1 and NINJ2 filaments. (A, B) Atomic models of two neighboring subunits in the NINJ1 (A) or NINJ2 (B) filament structures. In each case, one subunit is depicted in gray, while the other is color-coded for four named regions, with the protein sequence number of their boundaries shown in the bar above. AH, amphipathic helix; TM, transmembrane helix. (C-J) Side-by-side comparison of atomic interactions responsible for the assembly of the NINJ1 filament (C-F) with those for the NINJ2 filament (G-J). They are nearly identical between the two. (C) and (G) show hydrophobic interactions between the AH2 and the TM helices. For clarity, the hydrophobic sidechains of residues on the TM helices are not shown. (D) and (H) show interactions between polar residues on AH2 and those on the TM helices. The red dot in each panel represents a water molecule mediating the hydrogen bond interactions among the four residues. Green dashed lines depict hydrogen bonds. (E) and (I) highlight hydrophobic residues on AH1 that may contribute to the interaction with the TM helices beneath it. Again, hydrophobic sidechains on the TM helices are not shown for clarity. (F) and (J) show the abundant hydrogen bond (green dashed line) and salt bridge (red dashed line) interactions among residues on the AH1, the TM helices, and the AH1 of the next subunit. (K, L) Hydrophobicity plots for the outer surface (left panel) and the inner surface (right panel) of the NINJ1 filament (K) or the NINJ2 filament (L). In each case, three consecutive subunits are depicted. A hydrophobicity scale bar is placed at the bottom.

AH1 extended transversely to a neighboring subunit, across the lower-middle part of its TM1 and TM2, until it reached the AH1 of that subunit (Figure2A and 2B). In fact, the carboxyl group of Ser50 (Ser36) at the beginning of the second AH1 was even forming hydrogen bond with the amine group of His40 (His26) at the end of the first AH1 (Figure2F and 2J), just like any backbone hydrogen bond one would expect in a continuous alpha helix. The sidechain of His40 (His26) may also form hydrogen bond with the sidechain of Ser50 (Ser36) (Figure2F and 2J). The “last” (the most N-terminal in sequence) residue resolved on AH1, Asn39 (Asn25), had its sidechain turned around towards the alpha-helix backbone and then formed hydrogen bond with the backbone carboxyl of Leu52 (Leu38) in the kink region of the neighboring subunit (Figure2F and 2J). This turn-around also redirected the remaining N-terminal coil in the sequence (a.a. 1-38 in NINJ1, and 1-24 in NINJ2) towards the side of the filament and avoided clashing with the AH1 of the neighboring subunit. The two consecutive lysine residues on AH1, Lys44 (Lys30) and Lys45 (Lys31), were highly conserved in many NINJ sequences (FigureS4). In our structures, the sidechain of Lys44 (Lys30) extended downward and formed hydrogen bond with Asn119 (Asn105) on TM2 of the neighboring subunit, while the sidechain of Lys45 (Lys31) extended upward and formed salt bridge with Asp53 (Asp39) in its kink region (Figure2F and 2J). In addition to this vast interaction network formed by charged/polar residues, hydrophobic interactions may also played a minor role (Figure2E and 2I). Of particular note was the highly conserved Tyr41 (Tyr27), which inserted its aromatic sidechain into the cleft formed by the kink region, the TM1 and the TM2 of the neighboring subunit. Other hydrophobic interactions included those of Ala47 (Val33), Ala48 (Ala34), and Met51 (Met37) with the TM helices beneath them (Figure2E and 2I).

The remaining residues on AH1 and AH2 were more-or-less exposed on the surface and they were less conserved between NINJ1 and NINJ2. These included Ala42 (Ala28), Ser43 (Thr29), Ser46 (Ser32), Glu49 (Glu35), Ala55 (Ala41), Met58 (Met44), Ala59 (Ser45), Ser62 (Met48), Gln63 (Arg49), Lys65 (Lys51), Glu69 (Glu55), and Gln70 (Gln56). One would easily notice that the sidechains of these residues were limited to charged, polar, or only moderately hydrophobic ones. In fact, surface rendering of the protofilaments of NINJ1 and NINJ2 showed that their outer surface (the one facing the bulk solution) was highly hydrophilic (Figure2K and 2L), as hydrophobic residues on the TM1 and TM2 had all been concealed by the AH1 and AH2. In stark contrast but not unexpected, the inner surface (the one facing the lipid bilayer) of the protofilament was highly hydrophobic (Figure2K and 2L).

Another structural feature worth noting was the loop connecting AH2 to TM1. The density of this loop was very well resolved in both NINJ1 and NINJ2 filament structures, suggesting that it was highly rigid. This was somewhat unexpected for a Gly-Pro-Ser sequence. Modeling showed that the ring of Pro72 (Pro58) was stacked with the aromatic ring of Tyr77 (Tyr63) beneath it, fixing the conformation of the loop (FigureS5). It remains to be tested for any functional significance to have a more-or-less rigid linker at this position.

### How NINJ1 mediates PMR and why NINJ2 cannot

Although our NINJ1 and NINJ2 structures were both double-layer filaments, barely any protein-protein interactions could be observed between the two layers, suggesting that the double layer was likely formed during the purification process because of hydrophobic interactions mediated by the lipids associated with the “inner surface” of each layer. In a native membrane environment, a single-layer filament would be expected. Indeed, we had observed some single-layer NINJ1 filaments in our cryoEM images, which appeared to be much more flexible than the double-layer ones (FigureS6B). Forming a double layer likely had straightened and rigidified the filament, similar to the situation that double-stranded DNA has a persistence length over 60 times greater than that of single-stranded DNA^24,25^. In our preliminary purification trials of NINJ1, when the yield was low, we sometimes observed small circles of NINJ1 filaments (likely single-layer) (FigureS6A) in addition to the straight ones (double-layer), testifying the intrinsic lateral flexibility of single-layer NINJ1 filament.

The observation of a hydrophobic, lipid-binding inner surface and a hydrophilic, soluble outer surface of NINJ1 filament (Figure2K), in addition to the understanding of its lateral flexibility, intuitively pointed to two possible mechanisms for NINJ1 to mediate PMR. One mechanism is to form a hole on the plasma membrane like GSDMD, but in a larger size that would allow big molecules such as the DAMPs to pass through; the other mechanism is to wrap around a piece of membrane and directly “solubilize” it from the plasma membrane (Figure3A). As described in the previous section, we found that over-expressed NINJ1 was mostly associated with small membrane blebs in the media fraction instead of with the cell pellets. This observation strongly supports the second mechanism that NINJ1 is a “solubilizer” of the plasma membrane instead of a “hole puncher”. Interestingly, many membrane bleb-like particles could be directly observed in the negative-staining EM images of the purified NINJ1 sample (FigureS2E, FigureS6A), but not in the NINJ2 sample (FigureS2B). These were likely the smallest blebs that were somewhat resistant to detergent solubilization compared to large ones.

**Figure 3.**
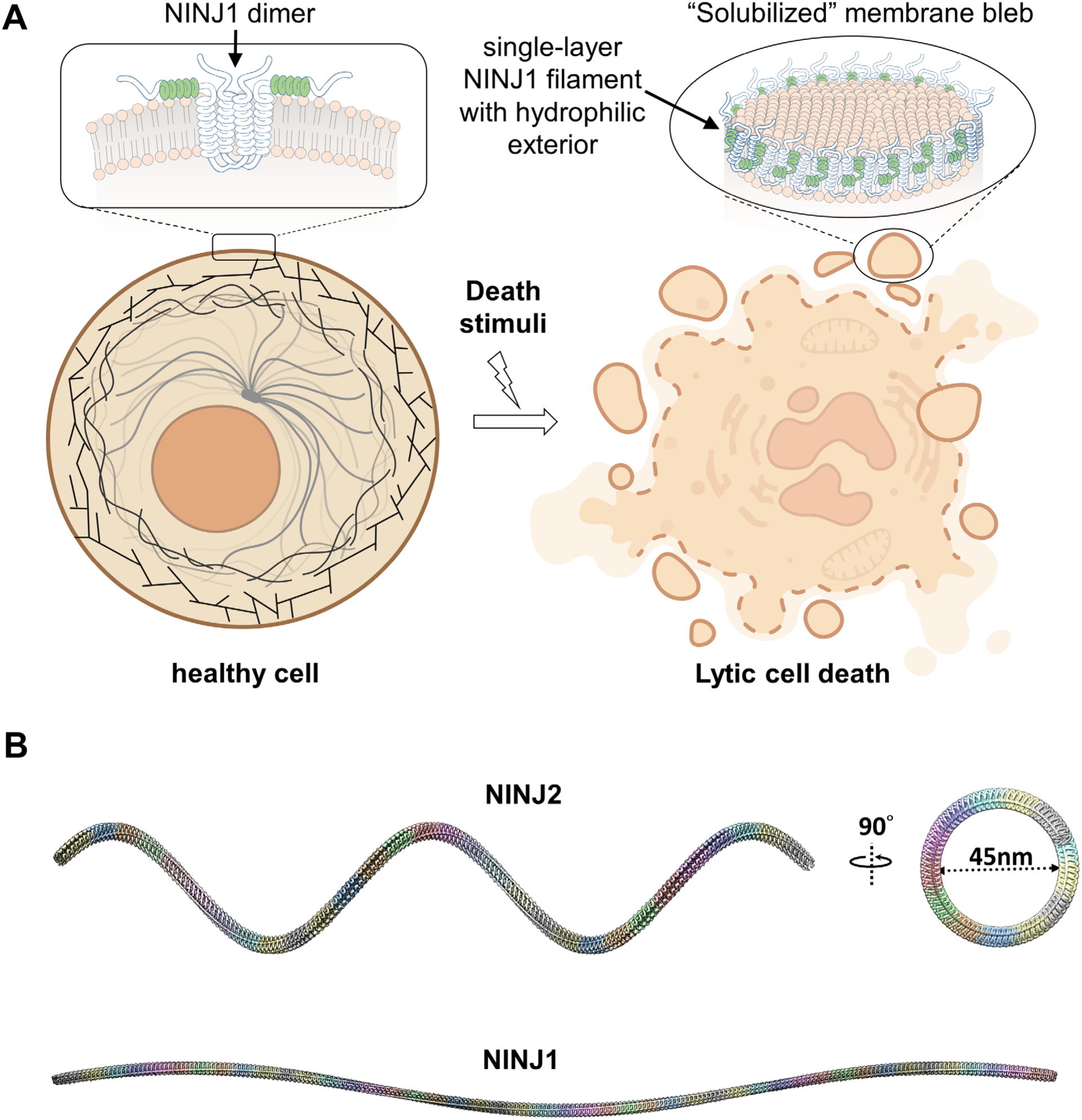
How NINJ1 mediates PMR and why NINJ2 cannot. (A) A proposed mechanism of NINJ1-mediated PMR. On the left, a healthy cell with intact plasma membrane (solid brown colored line) is shown. The magnified inset depicts a NINJ1 dimer (see discussion) with its amphipathic helices (highlighted in green) floating on the membrane surface. To the right, a dying cell with ruptured plasma membrane (dashed brown colored line) is depicted. Inset represents zoomed view of a “solubilized membrane bleb” wrapped at the edge by a single layer of NINJ1 filament. The NINJ1 filament has a hydrophobic inner surface interacting with the membrane and a hydrophilic outer surface rendering the membrane bleb “soluble”. (B) Virtually assembled long filaments of NINJ2 and NINJ1. Many copies of the cryoEM density map of NINJ2 or NINJ1 were fitted head-to-tail to extend the length of the filament. It indicated that NINJ2 filament could only wrap around a membrane tubule with a diameter of about 45 nm, explaining its incapability of mediating PMR in a regular cell, which is too flat for NINJ2 filament to assemble. In comparison, the NINJ1 filament is nearly straight.

With an understanding of how NINJ1 mediates PMR, we found that the incapability of NINJ2 in mediating PMR could be well explained by its intrinsically curved filament assembly. As discussed above, the single-layer NINJ1 filament has considerable lateral flexibility, which enables it to wrap around a membrane bleb and mediate PMR (Figure3A); however, vertical flexibility of the filament should be very limited, if not completely absent, as one can intuitively appreciate from the structure. The same should apply to the NINJ2 filament, which has a fixed curvature in the vertical direction. To quantify this curvature of the NINJ2 filament, we fitted many copies of the NINJ2 density map head-to-tail and obtained a spring-shaped virtual assembly (Figure3B). The diameter of this spring was measured to be about 45 nm (Figure3B). Imaginarily pressing this spring would be equivalent to applying the lateral flexibility of the NINJ2 filament. This virtual practice demonstrated that the only way for a single-layer NINJ2 filament to circle back to its starting point (note that this is still different from the circularization of the NINJ1 filament laterally) would be to assemble around a membrane tube or a vesicle that has a diameter of about 45 nm. Obviously, the plasma membrane of a regular cell with a diameter in the micron range would be too flat for the NINJ2 filament to wrap around and enclose.

### Lipid binding affects the curvature of NINJ1 and NINJ2 filament assembly and their capabilities of mediating PMR

As described above, the intermolecular interactions in NINJ1 and NINJ2 filaments are more or less the same, but the lipid densities resolved in the two filament structures were quite different. This prompted us to test whether lipid binding would affect the curvature of NINJ1 or NINJ2 filament as well as their capabilities of mediating PMR. At the inner leaflet, each subunit of NINJ2 was associated with three lipid densities (Figure4H and 4J), including one very prominent cholesterol-like density (position #3 in Figure4H) and two slightly weaker, fatty acid chain-like densities (position #1 and #2 in Figure4H). While in NINJ1, only one lipid density was associated with each protein subunit (Figure4B and 4D), at a position roughly corresponding to #1 in NINJ2 (Figure4B and 4H). This density was slightly planar and thus we modeled it as a cholesterol too (Figure4D and 4E). When NINJ1 and NINJ2 atomic models were superposed, the most significant structural difference was at the lower half of their TM2 (FigureS7A), where the cholesterol in NINJ2 binds (#3 in Figure4H). It could be regarded as if the NINJ2 TM2 was pushed by the very bulky cholesterol as it squeezed itself into the concave formed by two neighboring NINJ2 subunits, where it was crowded already by the other two lipid chains (Figure4H and 4I). This very subtle displacement of the TM2 makes the NINJ2 subunit more or less like an inverted trapezoid as compared to the rhomboid NINJ1, resulting in a gradual curving of the NINJ2 filament towards the inside of the cell (FigureS7B and S7C). The most important sequence variation that might be responsible for this discrepancy in cholesterol binding is probably Gln103 of NINJ2 versus Phe117 of NINJ1. Gln103 could form hydrogen bond with the hydroxyl group of an incoming cholesterol (Figure4K); while the bulky Phe117, together with Phe100 (corresponding to Val86 in NINJ2) from a neighboring subunit, sterically hindered the binding of a cholesterol at this place (Figure4B and 4C), as well as the binding of another lipid at a neighboring position corresponding to #2 in NINJ2 (Figure4B and 4H). Interestingly, this sequence variation between human NINJ1 and NINJ2 was conserved in mouse and rat too, even though mouse NINJ2 had a lysine residue (instead of glutamine) at this position (FigureS4), which could also form hydrogen bond with the hydroxyl group of cholesterol. Other conserved sequence variations, including Ile110 (Gly124 in NINJ1), Ala107 (Leu121) and Val79 (Gly93), might also contribute to the binding of the lipids or are necessary to reduce the steric hindrances (Figure4E and 4K).

**Figure 4.**
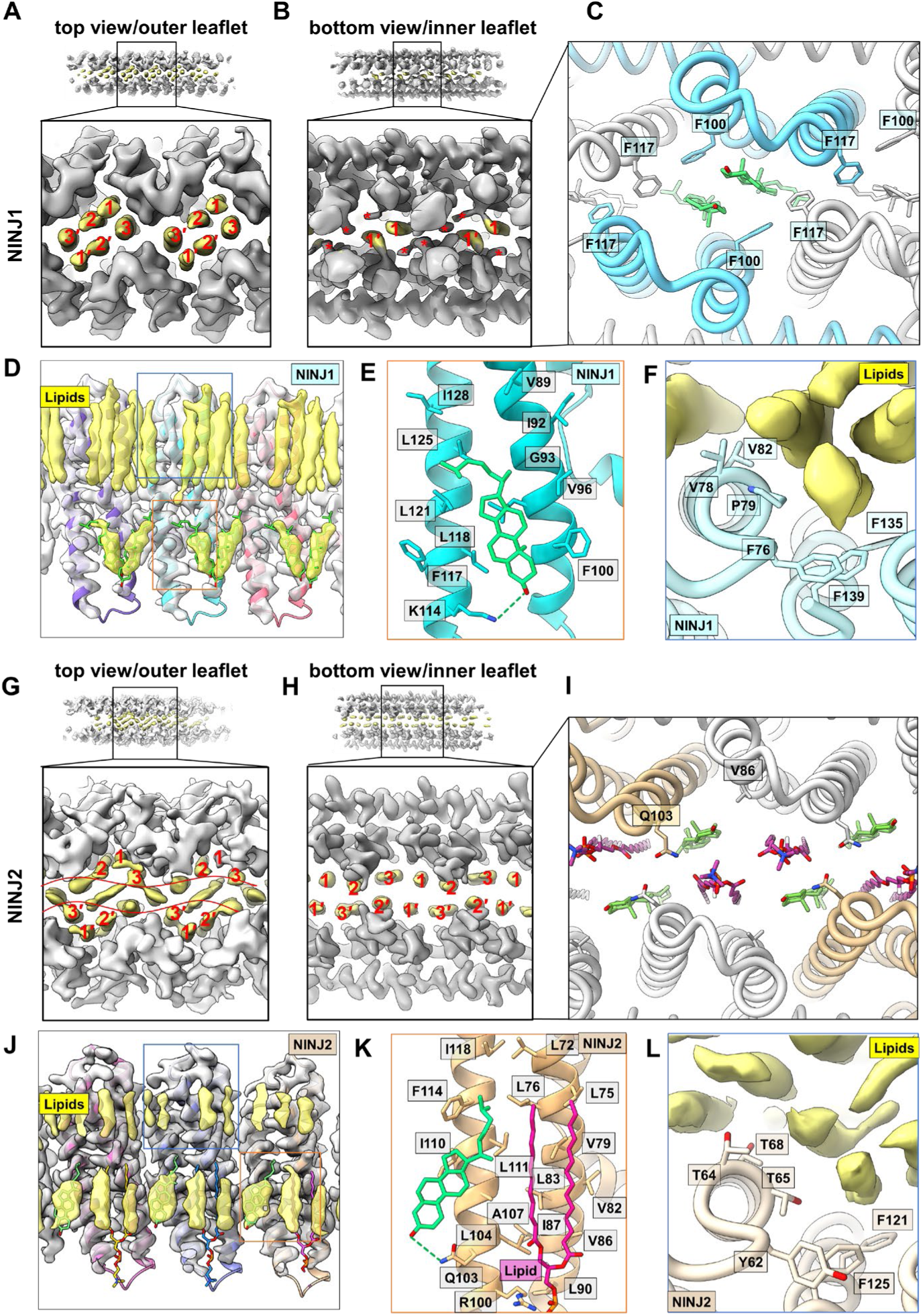
Differences in lipid binding between NINJ1 and NINJ2 filaments. (A) Zoom-in view of the NINJ1 filament from top. Lipid densities in the outer leaflet of the bilayer are highlighted in yellow and marked with numbers 1-3, or 1’-3’ on the other side at symmetrical positions. Roughly two repeating units of the filament are shown. (B) Zoom-in view of the NINJ1 filament from bottom to show the lipid densities in the inner leaflet of the bilayer, similar to (A). Note that densities marked with red “*” signs are sidechains of Phe100 or Phe117, not lipids. (C) Same view as (B), but shown with models and only for one repeating unit. (D) Exposed view of the lipid bilayer in NINJ1 filament. One layer of NINJ1 protein close to the viewer has been removed. The farther layer is semitransparent and fitted with atomic (colored ribbon). Densities in the inner leaflet are fitted with cholesterol models (green sticks). Densities in the outer leaflet are obviously lipids with aliphatic chains. They are not modeled because of unknown identity. (E) Atomic models showing interactions of the putative cholesterol (fitted in D) with the NINJ1 protein. This is the zoom-in view of the boxed region in D. (F) Protein-lipid interactions at the outer leaflet of the NINJ1 filament. The NINJ1 proteins are shown as ribbon models, with important sidechains discussed in the main text highlighted and shown as sticks. The un-modeled lipids with aliphatic chains are shown as yellow densities. This corresponds to a zoom-in view of the blue-boxed region in A. (G) Zoom-in view of the NINJ2 filament from top to show the lipid densities in the outer leaflet of the bilayer. Numberings of 1-3 and 1’-3’ are the same as A and can be match from one to the other in relative positions to the protein sidewall. The two wavy lines denote the extra layer of lipids in NINJ2 filament as compared to NINJ1. (H) Zoom-in view of the NINJ2 filament from bottom to show the lipid densities in the inner leaflet of the bilayer. To be compared with B. Position #3 is the critically important cholesterol discussed in the main text. (I) Same view as H with models. Gln103 and Val86 corresponding to Phe117 and Phe100 in C, respectively, are highlighted. (J) Exposed view of the lipid bilayer in NINJ2 filament. Same as D, but the lipids associated with the removed layer of NINJ2 protein are also removed for clarity. (K) Atomic models showing interactions of the cholesterol and lipid (fitted in J) with the NINJ2 protein. This is the zoom-in view of the boxed region in J. Note that the lipid density in the middle of the three and to the right of the cholesterol, modeled here as an aliphatic chain, can also be a cholesterol like in the case of NINJ1. The protein sidechains in this area are also consistent with a cholesterol binding. (L) Protein-lipid interactions at the outer leaflet of the NINJ2 filament. Same as F.

At the outer leaflet, as briefly described above, NINJ1-associated lipids were one-layer less than those in the NINJ2 filament (Figure4A and 4G), but their intensity was much higher (Figure4D and 4J), suggesting tighter association with the protein. We believe that four highly conserved (in mouse, human, and rat) sequence variations between NINJ1 and NINJ2 might have contributed to their different tightness in lipid association at the outer leaflet. These include Phe76, Val78, Pro79 and Val82 of NINJ1 versus Tyr62, Thr64, Thr65 and Thr68 of NINJ2, respectively (Figure4F and 4L). In NINJ1, the side chain of Phe76 was well aligned with those of Phe139 and Phe135, and the three were in turn stacked with the fatty acid chain of the associated lipid (Figure4F). In NINJ2, it seems that the replacement of phenylalanine with tyrosine (Tyr62) broke the good alignment with Phe125 (corresponding to Phe139 in NINJ1) and Phe121 (Phe135 in NINJ1) and affected the lipid binding (Figure4L). A similar stacking effect was observed between the lipid chain and Pro79 in NINJ1 (Figure4F), which was replaced by polar residue Thr65 in NINJ2 (Figure4L). Obviously, replacing the hydrophobic Val78 and Val82 in NINJ1 by polar residues Thr64 and Thr68, respectively, in NINJ2 would also affect the association of another lipid chain (Figure4F and 4L).

To test the functional significance of the above discussed sequence variations, we swapped the corresponding residues between NINJ1 and NINJ2 and made several mutants (Table S2). The M5 mutants had five mutations in residues involved in lipid binding at the inner leaflet, and M4 mutant had four mutations in residues involved in lipid binding at the outer leaflet, while M9 mutants combined all of them. It had been demonstrated before that the capability of NINJ1 to mediate PMR was correlated with its cytotoxicity when overexpressed in HEK293 cells, which could be measured by a “killing score”^3^. Following the same procedure, we compared the killing scores of NINJ1 and NINJ2 mutants with those of the wild type (WT). As expected, the NINJ1-M5 and NINJ1-M9 mutants had their killing scores dropped to nearly zero, at a similar level as NINJ2 WT (Figure5A), suggesting that lipid binding at the inner leaflet was indeed determinative for NINJ1’s capability of mediating PMR. The NINJ1-M4 mutant had a killing score about half of the WT (Figure5A), confirming that strong lipid binding at the outer leaflet (Figure4F) was also important for NINJ1’s functionality. Surprisingly, none of the NINJ2-M5 and NINJ2-M9 mutants gained capability of mediating PMR as we expected. A purification of the NINJ2-M9 mutant indicated that it completely failed to assemble into filament (Figure5C), explaining its incapability of mediating PMR. In retrospect, maybe some of the mutations in NINJ2-M5 and NINJ2-M9 was not compatible with other NINJ2-specific residues that we failed to take into account. For example, the sidechain of V86F might be clashing with the very bulky sidechain of a neighboring Trp99 (Ala113 in NINJ1) (FigureS4). To our satisfaction, the purified NINJ1-M9 mutant looked more like NINJ2 WT instead of NINJ1 WT, with obviously curved side views in negative staining EM (Figure5B and 5C). This confirmed that we had successfully phenocopied NINJ2 from NINJ1 by imposing cholesterol binding at the inner leaflet of the lipid bilayer, validating our interpretations of the sequence-function relationship in NINJ1 and NINJ2.

**Figure 5.**
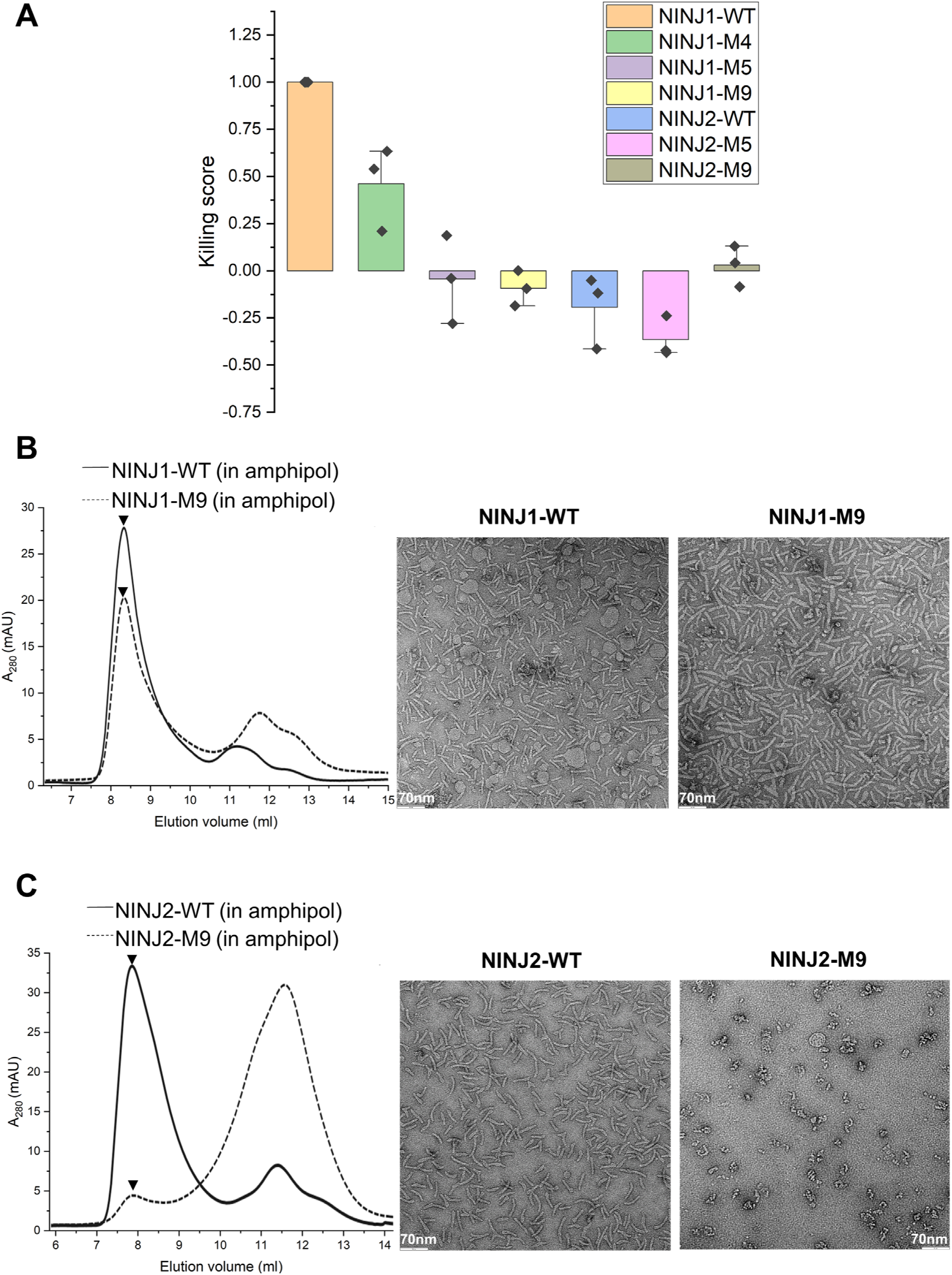
The role of lipid binding in the assembly and functionality of NINJ1 and NINJ2 filaments. (A) Cytotoxicity of NINJ1 or NINJ2 wild type (WT) and mutants. Cytotoxicity, correlated with NINJ1 or NINJ2’s capability of inducing PMR, is measured as killing score and plotted on the Y-axis. The height of each histogram corresponds to the mean killing score for the respective protein, and the bar at the top of each histogram indicates ±standard error of mean (SEM). The killing scores are calculated by normalizing the cytotoxicity of the mutant proteins to that of NINJ1-WT, as described in reference 3. (B) Comparing assembly of the NINJ1-M9 mutant with the NINJ1-WT. On the left, SEC profiles of NINJ1-WT and the NINJ1-M9 mutant purified and exchanged in amphipol.are plotted together. The Y-axis represents absorbance at 280 nm, while the X-axis represents elution volume in milliliters. Samples from the peak fractions (marked by arrowheads) are checked with negative-staining EM, and the micrographs are shown on the right. (C) Comparing assembly of the NINJ2-M9 mutant with the NINJ2-WT. Same as B. Note that the NINJ1-M9 mutant looks more like NINJ2-WT with curved filaments; while NINJ2-M9 mutant failed to oligomerize based on its SEC profile.

## Discussion

Here we use comparative cryoEM structural studies of NINJ1 and NINJ2 to understand not only how NINJ1 mediates PMR but also why its close paralog NINJ2 cannot. Our structures revealed a simple but elegant mechanism in which filamentous oligomerization of NINJ1 by conformational change of its amphipathic helix spontaneously triggers the segmentation and solubilization of a membrane bleb from the plasma membrane. While this filament assembly mechanism is well conserved in NINJ2, subtle differences in lipid binding lead to a curved NINJ2 filament that is defective in mediating PMR in regular cells. However, it is intriguing to think that NINJ2 might be able to mediate membrane segmentation in a highly curved or tubular membrane, e.g., the process of a glia cell or the axon of a neuron. The sequence variations between human NINJ1 and NINJ2 are mostly well conserved in mouse and rat (FigureS4), suggesting that NINJ2 might have evolved to have a specialized function instead of being a defective copy of NINJ1.

The importance of maintaining plasma membrane integrity cannot be overstated for any cell. So the action of NINJ1 must be tightly regulated. As NINJ1 is constitutively expressed on the plasma membrane, there must be a mechanism to stabilize it at the resting state and prevent it from spontaneously transitioning into the filament state. The key might be to prevent the AH from accessing the TM helices to form those stable interactions as observed in our filament structure. Among those interactions, quite a few are between charged/polar residues on the AH with those on the TM helices, e.g., Asn60 with Ser87, Gln91 and Asn133 (i.e., the “triad”), and Lys44 with Asn119. These polar residues, in addition to Asn120 and Thr123 next to Asn119, are highly conserved among all NINJ1 and NINJ2 sequences in human, mouse and rat (FigureS4). As polar residues do not like the hydrophobic environment in the membrane, they must be concealed. We propose that this conundrum can be solved by forming a homo-dimer of NINJ1, with the polar residues on TM helices facing each other at the dimer interface. This hypothesis is consistent with previous observation that endogenous NINJ1 in unstimulated bone marrow-derived macrophages migrated in SDS-PAGE as an approximately 16-kDa monomer, but in blue-native PAGE at approximately 40 kDa, likely as a dimer or trimer^3^. An Alphafold2 predicted dimer structure with the TM region only (the program tended to predict helix bundle between the TM region and the AH if the full-length sequence was used) seemed to be consistent with our hypothesis, with polar residues hidden inside and the hydrophobic residues facing outside (FigureS8A). Interestingly, a predicted trimer structure also made good sense chemically, but it would have a channel in the middle that seemed to be big enough to make the plasma membrane leaky (FigureS8B); therefore, we abandoned the trimer hypothesis. It was recently shown that glycine treatment protected cells against plasma membrane rupture by preventing NINJ1 oligomerization^26^. It seems possible in the context of our hypothesis that glycine may stabilize the NINJ1 dimer by mediating or strengthening the interactions among those polar residues at the dimer interface, thus slowing down or preventing the formation of NINJ1 filament. In particular, glycine might replace the water molecule associated with Ser87, Gln91 and Asn133 (Figure2D and 2H) and work better than the water in mediating their interactions at the dimer interface. Structural studies of NINJ1 at the resting state are needed to confirm these hypotheses.

If this dimeric structure proves to be true, it would be tempting to imagine the following scene of actions. As water flushes into a dying cell, through the GSDMD pores for example, the cell is ballooned and the plasma membrane is highly stretched. The surface tension of the membrane is pulling the NINJ1 dimer in two opposite directions perpendicular to their lipid-binding interface on the TM helices. This would at least weaken the interactions at the dimer interface, if it were too wild to think that the force might be strong enough to tear apart the dimer. The AH would now have access to the dimer interface and form those interactions needed for the assembly of the filament. As wild as it may seem, this mechanism of action could account for both the inertia of NINJ1 in a healthy cell and its quick action in a dying cell. For it to be possible, strong lipid binding is obviously the key.

At the time of manuscript preparation for this publication, a 3.8 Å resolution cryoEM structure of human NINJ1 expressed in *Escherichia coli* was reported by Degen et al.^27^. No data on NINJ2 was reported. A comparison of our atomic model of NINJ1 with theirs (PDB 8CQR) indicated that there was a three- or four-residue shift in registration for the amphipathic helices (AH1 and AH2 here, and α1 and α2 in the Degen et al. paper), while the modeling for the transmembrane helices was consistent between the two structures. As a result, the atomic interactions described in the Degen et al. paper were quite different from what was discussed here. As our NINJ1 structure has better resolution (2.75 Å vs 3.8 Å), we remain cautiously confident that our interpretation of the structure is more accurate. In addition, the mechanism of NINJ1-mediated PMR proposed in the Degen et al. paper was different from what we proposed in this paper.

## Acknowledgements

We would like to thank Brandon Miller for his help in the functional assay and insightful discussions, Dr. Kunpeng Li and Dr. Kyle Whiddon for their help with imaging at the Case Western Reserve University (CWRU) cryoEM facility. This work made use of the High Performance Computing Resource in the Core Facility for Advanced Research Computing at CWRU. This work was supported by the generous start-up funding from CWRU to XD.

## Author Contributions

X.D. and G.D. conceived the project; X.D. managed resources and supervised research; B.S., X.D. and Z.M. prepared samples, collected cryoEM data and determined the structures; B.S. and W.L. built atomic models; B.S. carried out functional assays; X.D., B.S. and W.L. analyzed the results and wrote the paper. All authors reviewed and approved the paper.

## Materials and Methods

### Expression and purification of NINJ1 and NINJ2 proteins

Mammalian expression constructs with pcDNA3.1(-) backbone and cDNA sequences for human NINJ1 (UniProtKB Q92982) or NINJ2 (UniProtKB Q9NZG7) protein with an N-terminal 3xFLAG tag and GSG linker were ordered from Genscript. For protein expression, about 1mg of purified plasmid was mixed with 3mL of 1mg/mL polyethylenimine (Polysciences) and transfected into 1L of Expi293F cells (Thermo Fisher) at cell density of ∼3.0 × 10^6^ cells/mL. Sodium butyrate was added at 10hr post transfection to a final concentration of 10mM to boost the protein expression level. The culture was harvested at 72hr post transfection for protein purification.

NINJ2 protein was purified from the cellular fraction of the culture. Cells were pelleted from the culture suspension by centrifuging at 6000×g for 20 minutes. The cell pellet was resuspended in TN buffer (50mM Tris-HCl pH 7.4, 150mM NaCl) supplemented with protease inhibitor cocktail (Roche), 1mM PMSF, 0.01% (w/v) NP40, 1% (w/v) glycerol, 10µg/mL DNase I (Roche), 5mM MgCl_2_, and 1mM CaCl_2_. The mixture was sonicated for 20 minutes at 40% amplitude with 5 sec on and 10 sec off cycles in a Qsonica sonicator, and then centrifuged at 4000xg for 20 minutes at 4℃ to remove large cell debris. The supernatant from this step was ultracentrifuged at 150,000×g for 60 minutes to pellet the membrane fraction. The membrane pellet was thoroughly resuspended in TN buffer using a homogenizer and then mixed with a pre-made solution of 10% (w/v) Dodecyl Maltoside (DDM) and 1% (w/v) Cholesteryl Hemisuccinate (CHS) (Anatrace) to a final concentration of 1% DDM and 0.1% CHS. The mixture was incubated for 3hr on a rotator at 4℃ for protein solubilization. The solubilized fraction was separated from the undissolved debris by another round of ultracentrifugation at 150,000×g for 30 minutes, and then incubated with pre-equilibrated M2 anti-FLAG affinity resin (Sigma-Aldrich) overnight on a rotator at 4℃. After draining on a gravity flow column, the resin was washed with 10 bed volumes of washing buffer (0.02% DDM, 0.002% CHS in TN buffer). The bound proteins were then reconstituted on-column into peptidisc^22^ or amphipol by incubation in 5ml of washing buffer supplemented with 10mg peptidisc peptide (Peptidisc Lab) or amphipol A8-35 (Anatrace) for 30 minutes. The mixture was then diluted stepwise (over a period of 90 minutes) using TN buffer to a total volume of ∼20ml to bring the detergent below its critical micelle concentration (CMC). The reconstitution buffer was then allowed to pass through the column followed by another round of washing with 10 bed volumes of TN buffer. The proteins were eluted from the column in 5mL TN buffer containing 0.2 mg/mL 3xFLAG peptide. The elution was concentrated using an Amicon centrifugal filter unit (Millipore) with a 30-kDa molecular weight cut-off and loaded onto a Superdex 200 Increase 10/300 GL column (Cytiva) for size-exclusion chromatography (SEC). The SEC fractions were checked with negative-staining electron microscopy on a Tecnai F20 microscope (Thermo Fisher). Selected fractions were combined and concentrated for cryoEM sample preparation.

NINJ1 purification followed the same procedure as NINJ2 except that the proteins were extracted from membrane blebs recovered from the culture media instead of from the cell pellet. Briefly, NINJ1-expressing Expi293F cell suspension harvested at 72hr post transfection was centrifuged at 6000xg for 20 minutes to remove the cells. The supernatant was collected and ultracentrifuged at 150,000xg for 40 minutes to pellet small membrane blebs released to the culture media due to NINJ1-mediated cell lysis (i.e., cytotoxicity). Six liters of culture were usually used for each batch of NINJ1 purification. The membrane fraction was then solubilized and processed following the same protocol detailed above for NINJ2 purification.

### Negative-staining electron microscopy

Carbon-coated copper grids were glow-discharged right before use. A 3.5-microliter aliquot of sample was applied to the carbon side of the grid and incubated for 1min. After the excess sample was blotted away with filter paper, the sample-loading side of the grid was flushed with three drops of deionized water followed by two drops of 2% (w/v) uranyl formate solution. A third drop of uranyl formate was kept on the grid for 1 min before blotting and drying of the grid. The grid was loaded into a Tecnai F20 microscope (Thermo Fisher) for imaging. Micrographs were recorded with a CCD camera usually at 50,000x magnification.

### CryoEM sample preparation, data collection, and data processing

The target protein was concentrated to ∼1mg/mL for cryoEM sample preparation. An aliquot of 3.5µL sample was applied to a 300-mesh Quantifoil R1.2/1.3 Cu grid pre-treated with glow-discharge. Excess liquid was blotted away using filter paper in a Vitrobot Mark IV machine (Thermo Fisher) with blotting force −5 and blotting time 3 seconds. The grid was plunge-frozen in liquid ethane.

CryoEM data collection was done with serialEM^28^ on a Titan Krios microscope (Thermo Fisher) equipped with Gatan BioQuantum K3 imaging filter and camera. A 10eV slit was used for the filter. Images were recorded at 130,000x magnification, corresponding to a pixel size of 0.33 Å/pix at super-resolution mode of the camera. A defocus range of −0.8µm to −1.5µm was set. A total dose of 50e^-^/Å^2^ for each exposure was fractionated into 50 frames. The first two frames of each movie stack were excluded in motion-correction. Frames were averaged with dose weighting^29^ and 2x binning, producing micrographs with a pixel size of 0.66Å/pix. To cope with strong preferred-orientation observed in both NINJ1 and NINJ2 samples, we also collected data with the sample tilted to 30° and combined it with untilted data.

CryoEM data processing was performed with cryoSPARC^30^ following the procedure for helical reconstruction, but no helical symmetry was applied throughout the data processing. Separation distance between segments in filament tracing was set to be 27.5Å (half of diameter, which was set to be 55Å), slightly larger than the 22Å separation distance of neighboring subunits as measured from initial reconstructions. To balance the number of particles from different views and alleviate the preferred-orientation artifact in reconstruction, only a portion of the highest-quality “top-view” particles were selected for reconstruction. Statistics of the datasets and structures are detailed in Table S1.

### Model building and refinement

Alphafold2-predicted human NINJ1 or NINJ2 atomic models were downloaded from the Alpha Fold Protein Structure Database^31^ and fitted into the density maps with UCSF Chimera^32^ to facilitate initial residue registration. The models were manually adjusted in Coot^33^ and iteratively refined with Phenix real-space refinement^34^.

### Data availability

CryoEM density maps and the associated atomic models for human NINJ1 and NINJ2 have been deposited to the Electron Microscopy Data Bank (EMDB) and the Protein Data Bank (PDB). The accession codes are EMD-40905 and 8SZA for NINJ1, and EMD-40907 and 8SZB for NINJ2.

### Cytotoxicity assay

For cytotoxicity assay, we constructed lentiviruses expressing each protein of interest to avoid background cytotoxicity associated with lipid-based transfection reagents. Firstly, lentivirus transfer constructs were prepared by cloning the NINJ1 and NINJ2 WT/mutant ORFs from the pcDNA3.1(-) vectors to pLL3.7 vector under the CMV promoter. These constructs, along with lentiviral packaging (psPAX2) and envelope (pMD2.G) plasmids, were transfected at equimolar ratios into adherent HEK293T cells using Lipofectamine 3000 reagent (Thermo Fisher). The pLL3.7 plasmid was a gift from Luk Parijs (Addgene plasmid #11795)^35^; pMD2.G and psPAX2 was a gift from Didier Trono (Addgene plasmid #12259 and #12260, respectively). Lentiviruses were harvested 72hr post transfection, filtered, and concentrated by centrifugation. A control batch of GFP-expressing lentiviruses were always included as control to visually inspect the quality and titer of lentivirus preparations using a fluorescence microscope. The lentiviruses were then used at an MOI of 5 to infect fresh HEK293T cells at ∼80% confluency for the cytotoxicity assay. The assay was performed at 16hr post infection using the CellTox Green Cytotoxicity Assay Kit (Promega). Fluorescence readings were recorded as triplicates using a Tecan Infinite M1000 Quadruple Monochromator Microplate Reader. The experiment was repeated three times in total. Pooled data were analyzed in Excel (Microsoft) and the graph was prepared with Origin (OriginLab).

---

**Table S1.**
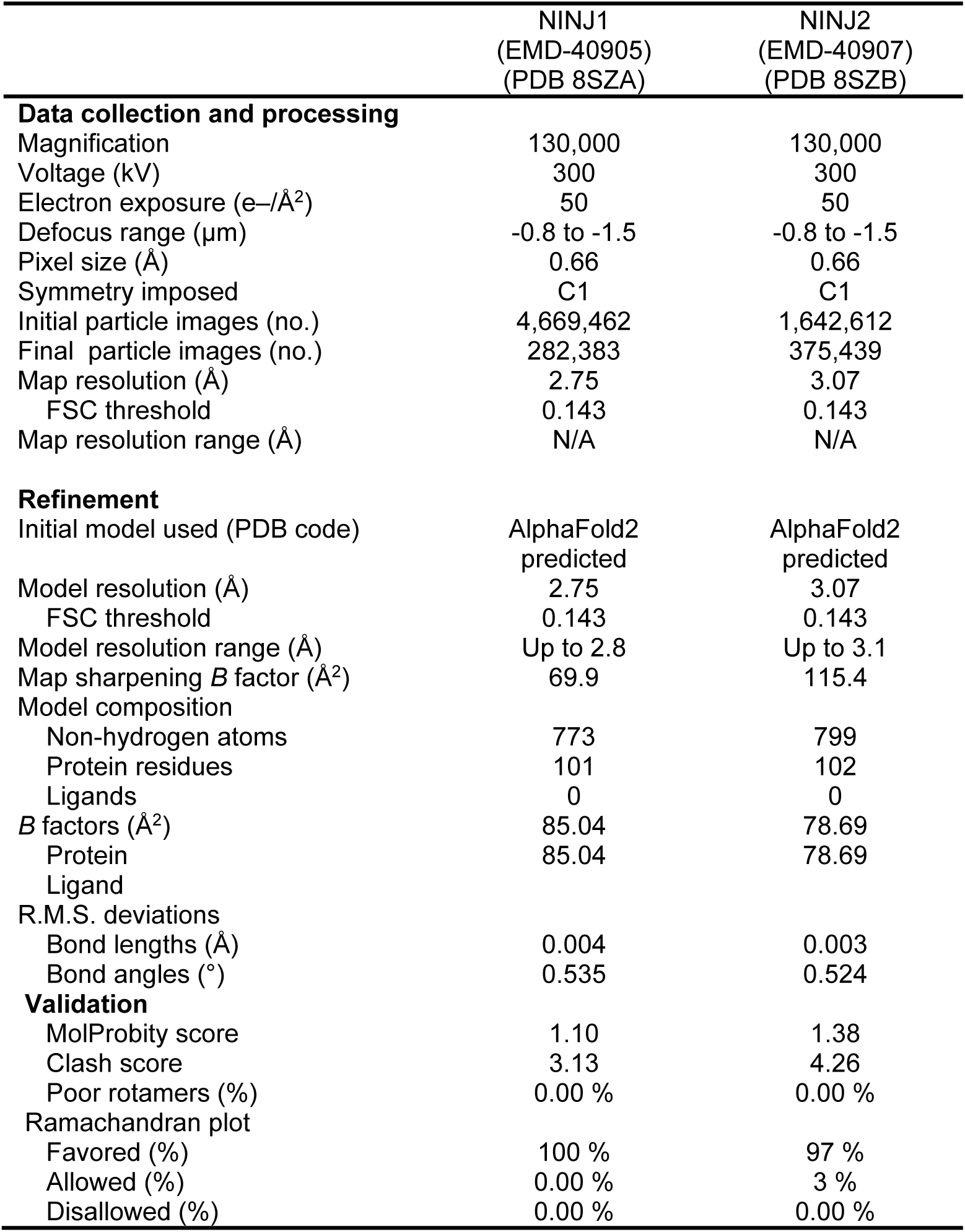
CryoEM Data Collection, Refinement and Validation statistics.

|  | NINJ1<br>(EMD-40905)<br>(PDB 8SZA) | NINJ2<br>(EMD-40907)<br>(PDB 8SZB) |
| --- | --- | --- |
| **Data collection and processing** |  |  |
| Magnification | 130,000 | 130,000 |
| Voltage (kV) | 300 | 300 |
| Electron exposure (e-/Å<sup>2</sup>) | 50 | 50 |
| Defocus range (µm) | -0.8 to -1.5 | -0.8 to -1.5 |
| Pixel size (Å) | 0.66 | 0.66 |
| Symmetry imposed | C1 | C1 |
| Initial particle images (no.) | 4,669,462 | 1,642,612 |
| Final particle images (no.) | 282,383 | 375,439 |
| Map resolution (Å) | 2.75 | 3.07 |
| FSC threshold | 0.143 | 0.143 |
| Map resolution range (Å) | N/A | N/A |
| **Refinement** |  |  |
| Initial model used (PDB code) | AlphaFold2<br>predicted | AlphaFold2<br>predicted |
| Model resolution (Å) | 2.75 | 3.07 |
| FSC threshold | 0.143 | 0.143 |
| Model resolution range (Å) | Up to 2.8 | Up to 3.1 |
| Map sharpening *B* factor (Å<sup>2</sup>) | 69.9 | 115.4 |
| Model composition |  |  |
| Non-hydrogen atoms | 773 | 799 |
| Protein residues | 101 | 102 |
| Ligands | 0 | 0 |
| *B* factors (Å<sup>2</sup>) | 85.04 | 78.69 |
| Protein | 85.04 | 78.69 |
| Ligand |  |  |
| R.M.S. deviations |  |  |
| Bond lengths (Å) | 0.004 | 0.003 |
| Bond angles (°) | 0.535 | 0.524 |
| **Validation** |  |  |
| MolProbity score | 1.10 | 1.38 |
| Clash score | 3.13 | 4.26 |
| Poor rotamers (%) | 0.00 % | 0.00 % |
| Ramachandran plot |  |  |
| Favored (%) | 100 % | 97 % |
| Allowed (%) | 0.00 % | 3 % |
| Disallowed (%) | 0.00 % | 0.00 % |

**Table S2.** Human NINJ1 and NINJ2 Mutants.

| NINJ1-WT | NINJ2-WT | NINJ1-M5 | NINJ1-M4 | NINJ1-M9 | NINJ2-M5 | NINJ2-M9 |
| --- | --- | --- | --- | --- | --- | --- |
| G93 | V79 | G93V |  | G93V | V79G | V79G |
| F100 | V86 | F100V |  | F100V | V86F | V86F |
| F117 | Q103 | F117Q |  | F117Q | Q103F | Q103F |
| L121 | A107 | L121A |  | L121A | A107L | A107L |
| G124 | I110 | G124I |  | G124I | I110G | I110G |
| F76 | Y62 |  | F76Y | F76Y |  | Y62F |
| V78 | T64 |  | V78T | V78T |  | T64V |
| P79 | T65 |  | P79T | P79T |  | T65P |
| V82 | T68 |  | V82T | V82T |  | T68V |

**Figure S1.**
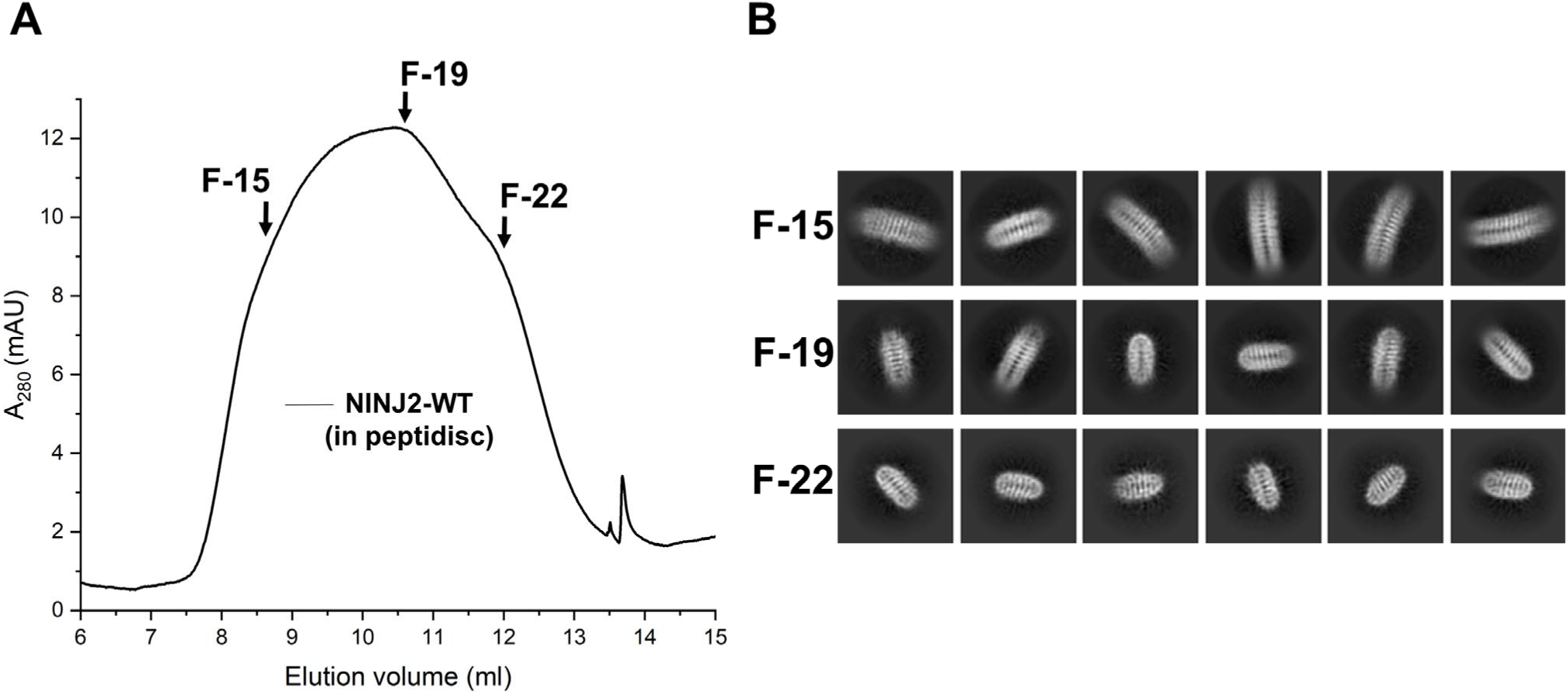
NINJ2 forms linear oligomers, related to Figure 1. (A) Size-exclusion chromatography (SEC) profile of purified human NINJ2 protein reconstituted in peptidisc. The X-axis represents elution volume in milliliters, while the Y-axis represents absorbance at 280 nm. Three representative peak fractions, marked with arrows, were taken for cryoEM checking. (B) Representative 2D class averages of cryoEM imaging for those three fractions.

**Figure S2.**
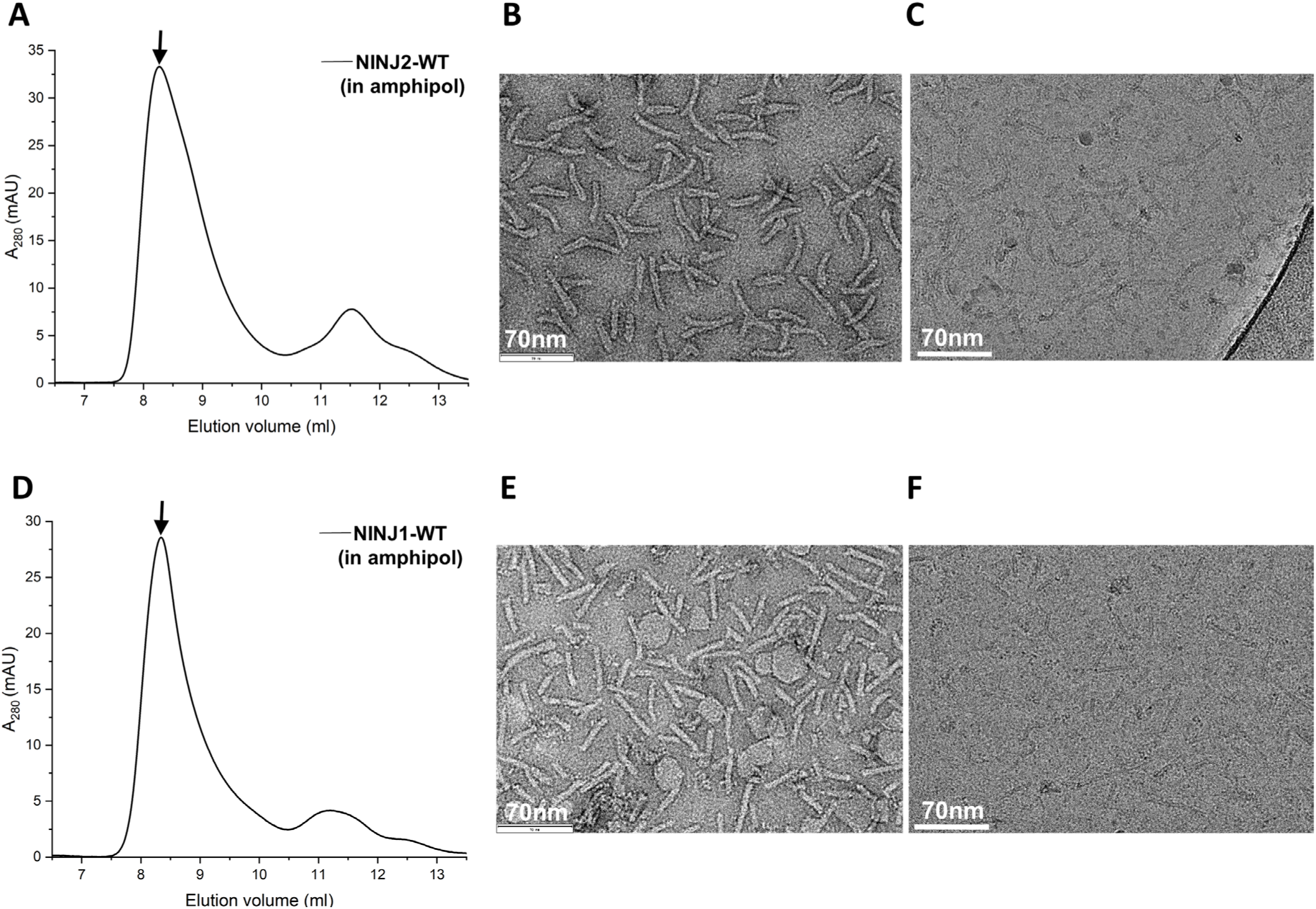
Sample preparation of homogeneous NINJ2 and NINJ1 filaments for high-resolution cryoEM structure determination, related to Figure 1. (A) The SEC profile of human NINJ2 purified and exchanged in amphipol A8-35. The X-axis represents elution volume in milliliters, while the Y-axis represents absorbance at 280 nm. The peak fraction used in panels B and C is indicated with an arrow. (B, C) A representative negative-staining EM (B) and cryoEM (C) micrograph of the purified NINJ2 sample. (D) The SEC profile of human NINJ1 purified and exchanged in amphipol A8-35. (E, F) A representative negative-staining EM (E) and cryoEM (F) micrograph of the purified NINJ1 sample.

**Figure S3.**
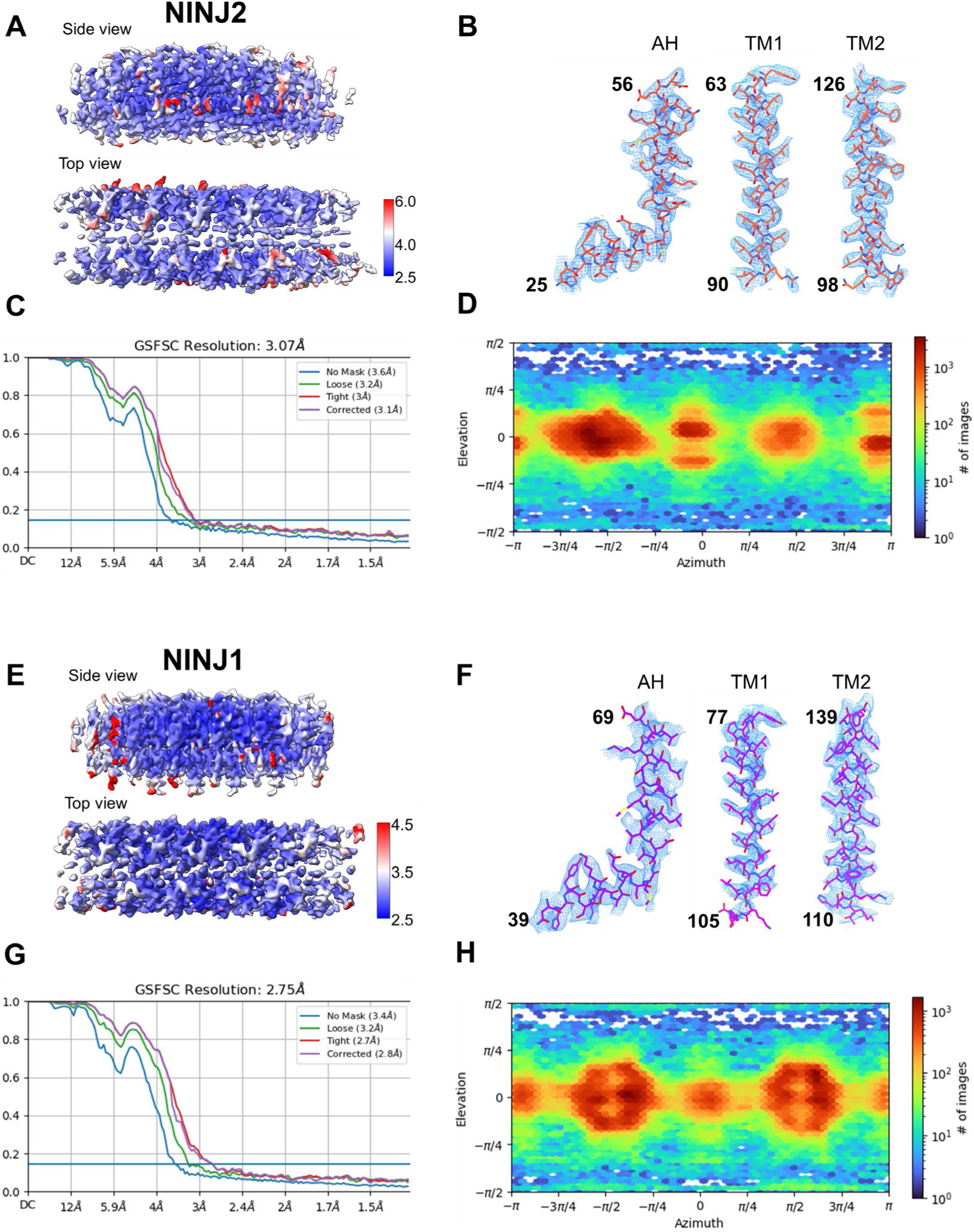
Quality assessment of the cryoEM structures of the NINJ2 and NINJ1 filaments, related to Figure 1. (A) Local resolution of the cryoEM density map of the NINJ2 filament. Side and top views of the NINJ2 filament were surface colored as per local resolution values. A scale bar is presented to the right. (B) High resolution features of the NINJ2 cryoEM structure. CryoEM densities are shown as blue-colored mesh, while the fitted model as orange-colored sticks. Residue numbers are labeled for the N- and C-termini of the models. From left to right, the amphipathic helix, and the transmembrane helices TM1 and TM2 are displayed. (C) Gold standard Fourier shell correlation (GSFSC) plot of the NINJ2 cryoEM structure. (D) Orientation distribution plot for particles (n = 375,439) used in the final reconstruction of the density map of the NINJ2 filament. (E) Local resolution of the cryoEM density map of the NINJ1 filament. Side and top views of the NINJ1 filament were surface colored as per local resolution values. A scale bar is presented to the right. (F) High resolution features of the NINJ1 cryoEM structure. CryoEM densities are shown as blue-colored mesh, while the fitted model as purple-colored sticks. Residue numbers are labeled for the N- and C-termini of the models. From left to right, the amphipathic helix, and the transmembrane helices TM1 and TM2 are displayed. (G) Gold standard Fourier shell correlation (GSFSC) plot of the NINJ1 cryoEM structure. (H) Orientation distribution plot for particles (n = 282,383) used in the final reconstruction of the density map of the NINJ1 filament.

**Figure S4.**
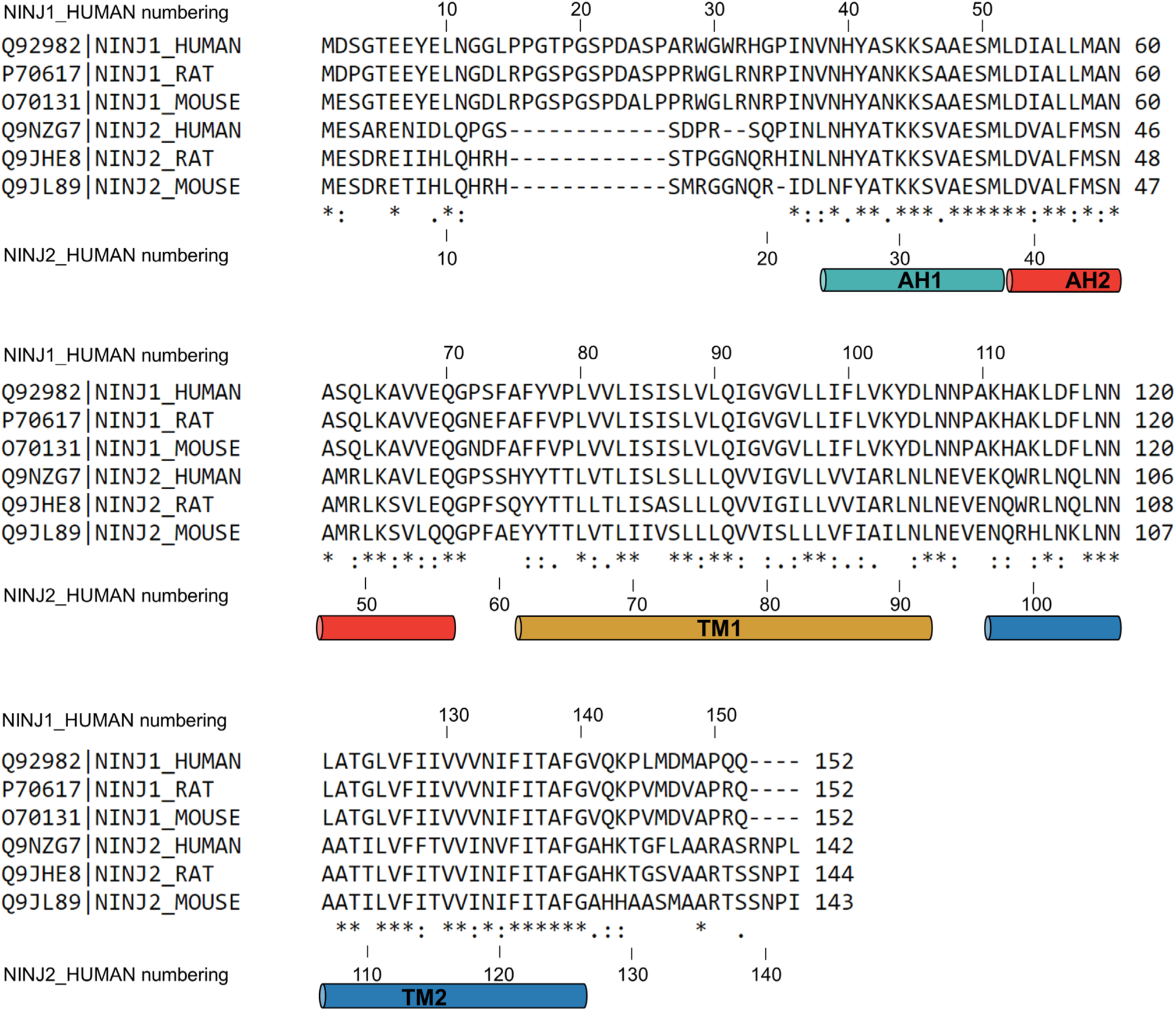
Sequence variations between NINJ1 and NINJ2 are highly conserved among different species, related to Figure 2 and Figure 4. Multiple-sequence alignment is shown for human, rat, and mouse NINJ1 and NINJ2 proteins. UnitProt ID was indicated for each sequence used for alignment. Amino acid sequence numbering for human NINJ1 is shown above the alignment, and that of human NINJ2 shown below, for easy location of each residue of interest. Secondary structures of the amphipathic helices (AH1 and AH2) and transmembrane helices (TM1 and TM2) discussed in the main text are annotated.

**Figure S5.**
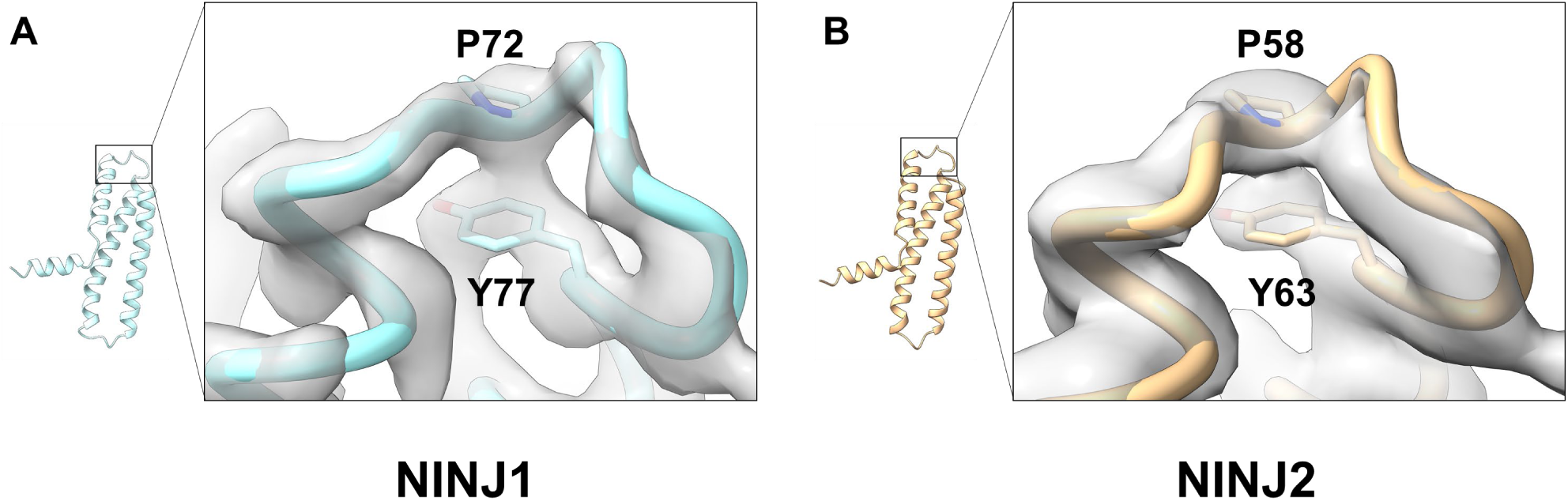
Highly rigid loop structures in NINJ1 and NINJ2, related to Figure 2. Structures of the loop region linking amphipathic helix AH2 and transmembrane helix TM1 are shown for NINJ1 (A) and NINJ2 (B) respectively. Location of the loop in the NINJ1 or NINJ2 subunit structure is highlighted by the box. In each case, the density map of the loop region is shown as semitransparent surface, with the atomic model (ribbon) fitted in. Stacking sidechains of the proline and tyrosine residues leading to the very rigid loop structures are highlighted with sticks.

**Figure S6.**
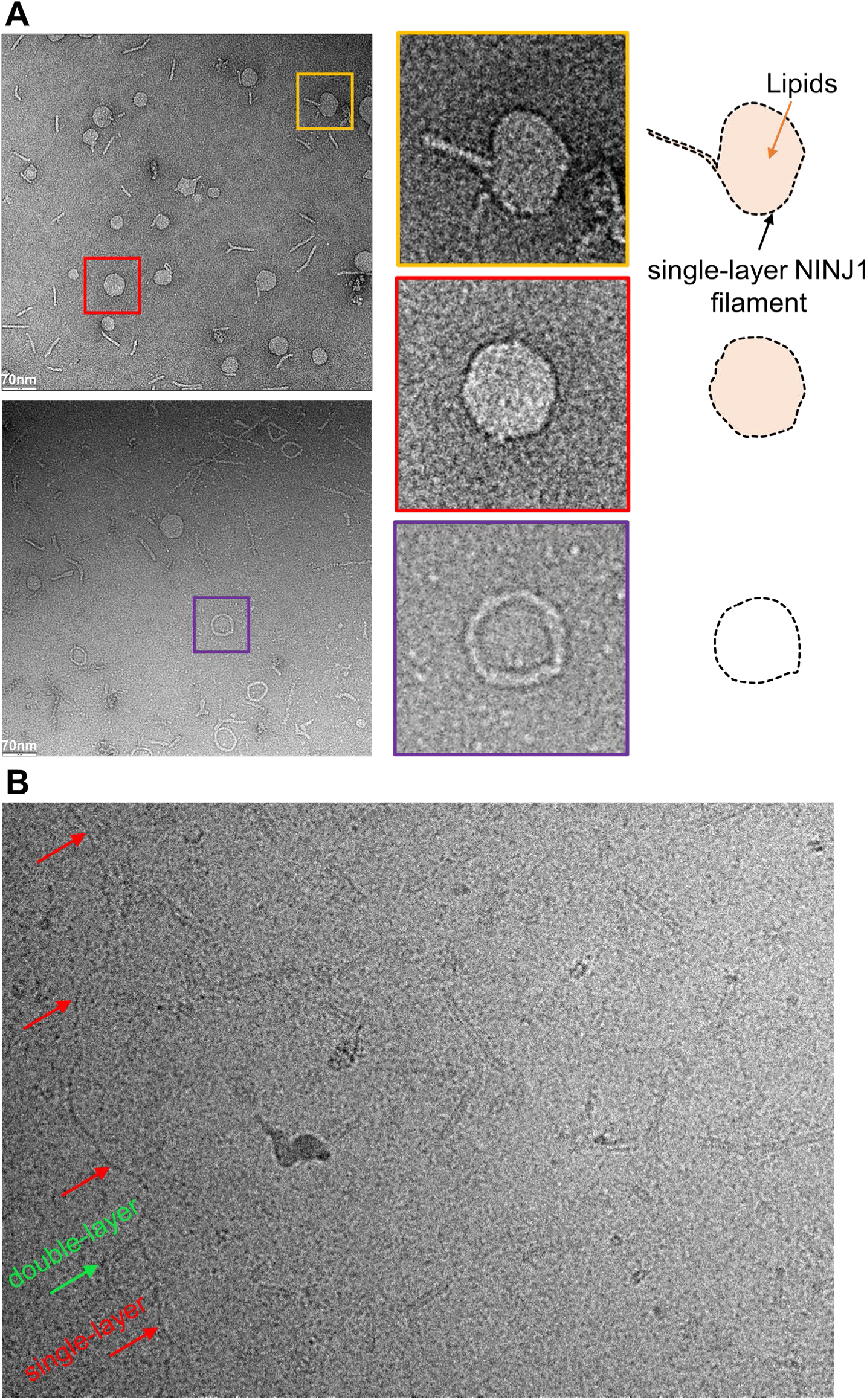
Single-layer NINJ1 filament is flexible and can wrap around membrane blebs, related to Figure 3. (A) Observation of membrane bleb-like (top left), or circle-like (bottom left) particles in negative-staining EM micrographs of purified NINJ1 samples. Zoom-in version of particles highlighted with boxes in the left are shown in the middle, with schematic interpretation on the right. (B) A long, flexible single-layer NINJ1 filament spotted in a micrograph of the cryoEM dataset. The red arrows point to single-layer sections of the filament; a green arrow points to a short double-layer segment of the filament. Most other particles are relatively short and straight double-layer filaments that we used for cryoEM structure determination.

**Figure S7.**
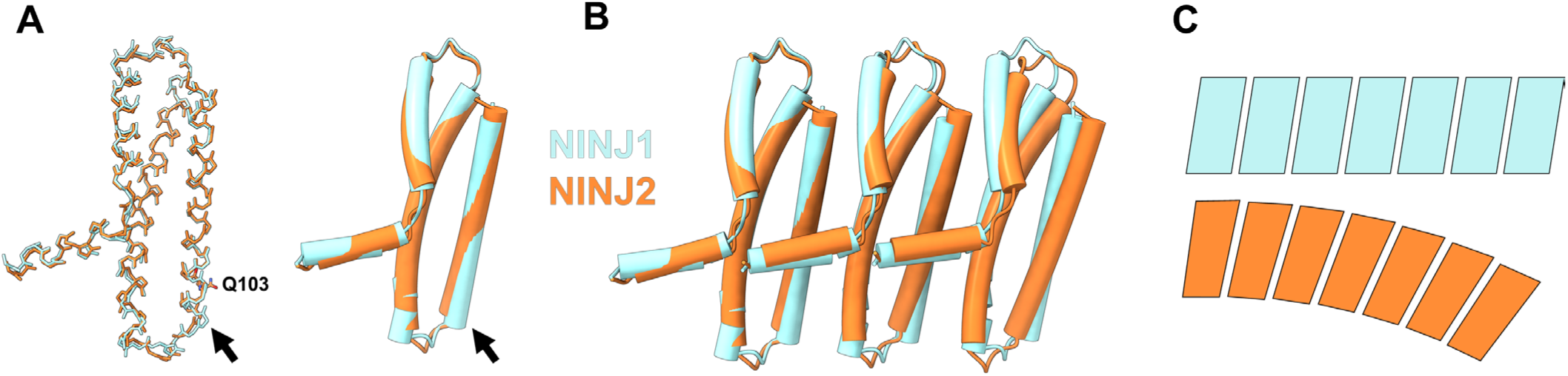
Structural basis for the curving of the NINJ2 filament as compared to the NINJ1 filament, related to Figure 4. (A) Comparing NINJ1 and NINJ2 subunit structures. Atomic models of NINJ1 (cyan) and NINJ2 (orange) are superposed and shown as backbones (left) or cylinders (right). The arrows point to where the two structures have the most “significant” difference. This happens to be where the cholesterol in NINJ2 binds. The determinative cholesterol-binding residue Gln103 of NINJ2 is highlighted. Note the narrowing in distance between the two transmembrane helices in NINJ2 as compared to NINJ1 because of this structural difference. (B) Comparing three consecutive subunits of NINJ1 and NINJ2. It shows how propagation and accumulation of the subtle structural difference depicted in A lead to the gradual curving of the NINJ2 filament as compared to the NINJ1 filament. (C) A schematic representation of (B) with more subunits.

**Figure S8.**
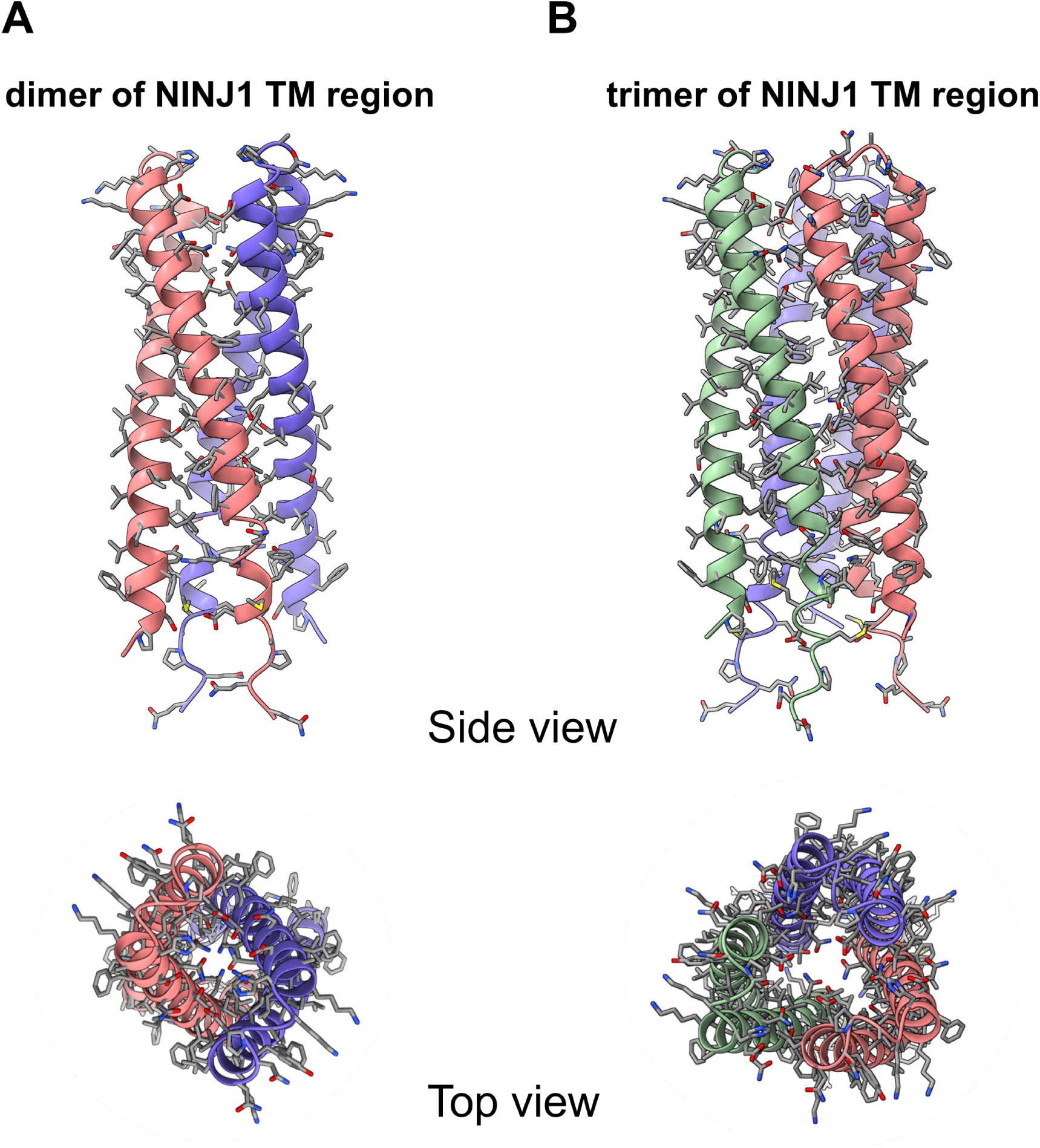
Possible conformation of human NINJ1 at resting state, related to Figure 3. (A) Alphafold2 predicted dimer structure of the NINJ1 transmembrane region. (B) Alphafold2 predicted trimer structure of the NINJ1 transmembrane region. Note that most polar residues are hidden in the dimer or trimer interface, away from the membrane environment. There seems to be a channel in the predicted trimer structure, which would make the membrane leaky.

